# Computational Modeling and Visualization of Ischemic Effects on an Advanced Purkinje Network

**DOI:** 10.1101/2025.06.21.660878

**Authors:** Laiba Khan, Maham Khan, Rizwan Ahmed, Mahmood Ahmad, Joanne Lac

## Abstract

The cardiac Purkinje network plays a vital role in the heart’s electrical conduction system, ensuring efficient and synchronized ventricular contraction. When impaired—particularly by ischemia—it can trigger life-threatening arrhythmias. In this study, we present an advanced computational model of the Purkinje network that integrates cell-level heterogeneity, spatial organization, and localized ischemic zones with customizable severity gradients. Developed in Python using open-source libraries (NumPy, matplotlib, pandas, seaborn), the model generates rich visualizations of network structure, conduction velocities, ischemia severity, and electrophysiological parameters. Simulations demonstrate how ischemia alters conduction and refractoriness in a spatially dependent manner, providing insights into arrhythmogenic risk. This modeling framework can advance understanding of cardiac conduction under pathological conditions and support therapeutic development.

## 1. Introduction

The heart’s electrical system relies on precise timing and coordination. Central to this system is the Purkinje network—a branching system of specialized fibers that rapidly conduct impulses through the ventricles. Myocardial ischemia, which restricts blood supply, can impair this network, disrupting normal impulse propagation and leading to arrhythmias such as ventricular tachycardia or fibrillation [1,5].

Computational modeling offers a powerful, non-invasive approach to investigate these mechanisms. While existing Purkinje fiber models have provided important insights [1–3], many lack integration of spatial anatomy, cellular heterogeneity, and detailed ischemic effects. To address these gaps, we developed a comprehensive computational framework simulating and visualizing ischemia-induced electrophysiological disturbances in a 3D Purkinje network.

## 2. Methods

### 2.1. Purkinje Network Model

We implemented the AdvancedPurkinjeNetwork class in Python to simulate a network of n Purkinje cells. Each cell includes spatial coordinates (x, y, z), randomly distributed within a bounded 3D volume to simulate biological dispersion. Cells are categorized as bundle_branch, fascicle, or purkinje to reflect histological diversity. A Boolean pacemaker flag allows some cells to exhibit spontaneous firing. Key electrophysiological parameters include conduction velocity, refractory period, and ATP level. To simulate pathology, each cell carries an is_ischemic flag and a continuous ischemia_severity score (0.0–1.0). Network connectivity is represented by a binary adjacency matrix, currently randomized but designed for future integration of anatomical branching data [4].

### 2.2. Simulating Ischemia

We developed an algorithm to define ischemic regions by specifying a center, radius, and severity. The Euclidean distance from each cell to the ischemic core determines individual severity, applied via linear or exponential gradients. Linear gradients model gradual decline, while exponential gradients simulate more abrupt severity transitions, consistent with ischemic tissue decay [5–7].

### 2.3. Visualization and Simulation

The ComprehensivePurkinjeSimulator class generates detailed visualizations to illustrate how ischemia impacts the Purkinje network. It includes 3D spatial plots of the cell layout, conduction velocity histograms, ischemia severity maps, and heatmaps of network connectivity. Additionally, it visualizes refractory period distributions and 2D projections of the cell positions. These individual components are integrated using the create_advanced_visualizations() function, which compiles them into a cohesive, publication-quality figure. This comprehensive visualization approach enhances interpretation of both structural organization and functional alterations within the network.

## 3. Results

Simulations with 500 Purkinje cells and two ischemic zones (one linear, one exponential) revealed the following patterns:

- 3D network views showed an interconnected, spatially dispersed layout, approximating biological architecture.
- Conduction velocities exhibited heterogeneous distributions, consistent with experimental variability [2].
- Ischemia severity maps accurately reflected the prescribed gradient profiles.
- Connectivity heatmaps revealed randomized but dense intercellular coupling.
- Refractory periods varied significantly across cells, supporting the modeling of reentrant conditions.
- 2D projections aided spatial interpretation of clustering and ischemic overlap.

## 4. Discussion

This modeling approach effectively captures how ischemia alters key electrophysiological parameters, such as conduction velocity and refractory period, while enabling direct visual correlation between spatial structure and functional disruption. Notably, areas with high ischemia severity often align with slowed conduction or shortened refractoriness—both recognized as precursors to arrhythmia [5–7]. While the current model uses randomized cell properties and connectivity, it is designed for future extensibility. Planned enhancements include integrating anatomically accurate branching patterns based on imaging data [4], implementing detailed electrophysiological models like Hodgkin-Huxley or Rudy-Li [1, 2], simulating action potential propagation with dynamic feedback, and exploring pharmacological interventions to assess therapeutic effects [3, 7].

Limitations of this model include its current focus on static representation of ischemia and cell properties. Dynamic simulations of impulse propagation, feedback mechanisms, and the temporal evolution of ischemic injury are not yet integrated. The cell model is simplified, and detailed ion channel kinetics are not considered.

Limitations: This model currently focuses on static representation of ischemia and cell properties. Dynamic simulations of impulse propagation, feedback mechanisms, and the temporal evolution of ischemic injury are not yet integrated. The cell model is simplified, and detailed ion channel kinetics are not considered.

Future Work:Future work will focus on implementing dynamic action potential propagation, developing anatomically informed connectivity models, and integrating biophysically detailed Purkinje cell models. We also aim to simulate pharmacological interventions and validate model predictions using experimental data from in vitro or in vivo studies.

## 5. Conclusion

We have developed a computational framework that models and visualizes how ischemia affects the Purkinje network. It integrates cell-level variability, customizable ischemic regions, and rich visual outputs, all within a Python-based ecosystem. The model provides a clear, visual, and quantitative understanding of how ischemic damage disrupts electrical conduction and may contribute to arrhythmia. With future enhancements, it holds strong potential for in silico exploration of cardiac pathology and intervention design.

## Grant Information

The author(s) declared that no grants were involved in supporting this work.

## Acknowledgments

None

## Figure Captions

**Figure 1A:**
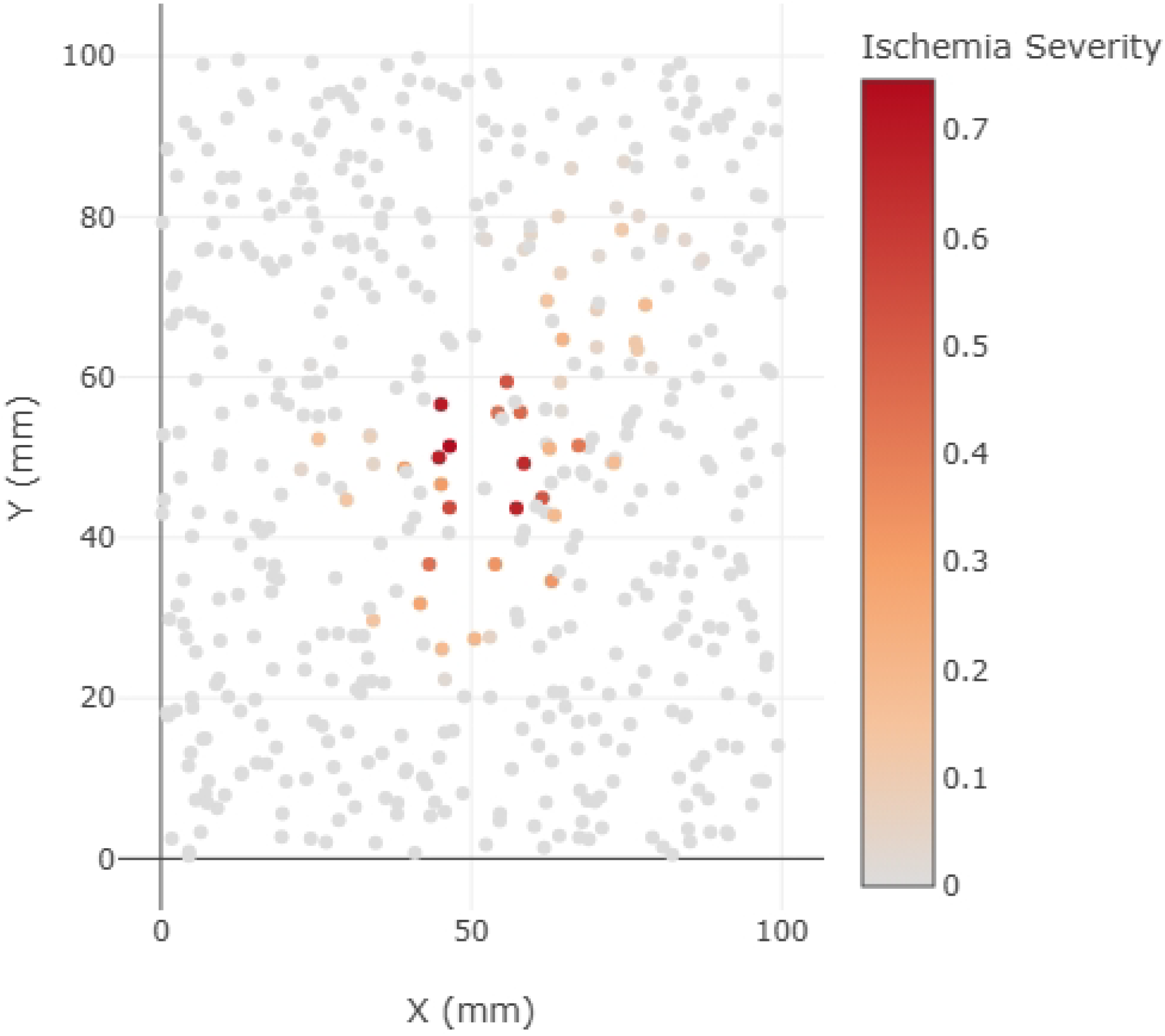
Three-dimensional visualization of the Purkinje network containing 500 cells randomly distributed within a cubic spatial domain. Cell positions reflect spatial dispersion approximating Purkinje fiber branching within the ventricular myocardium.

**Figure 1B:**
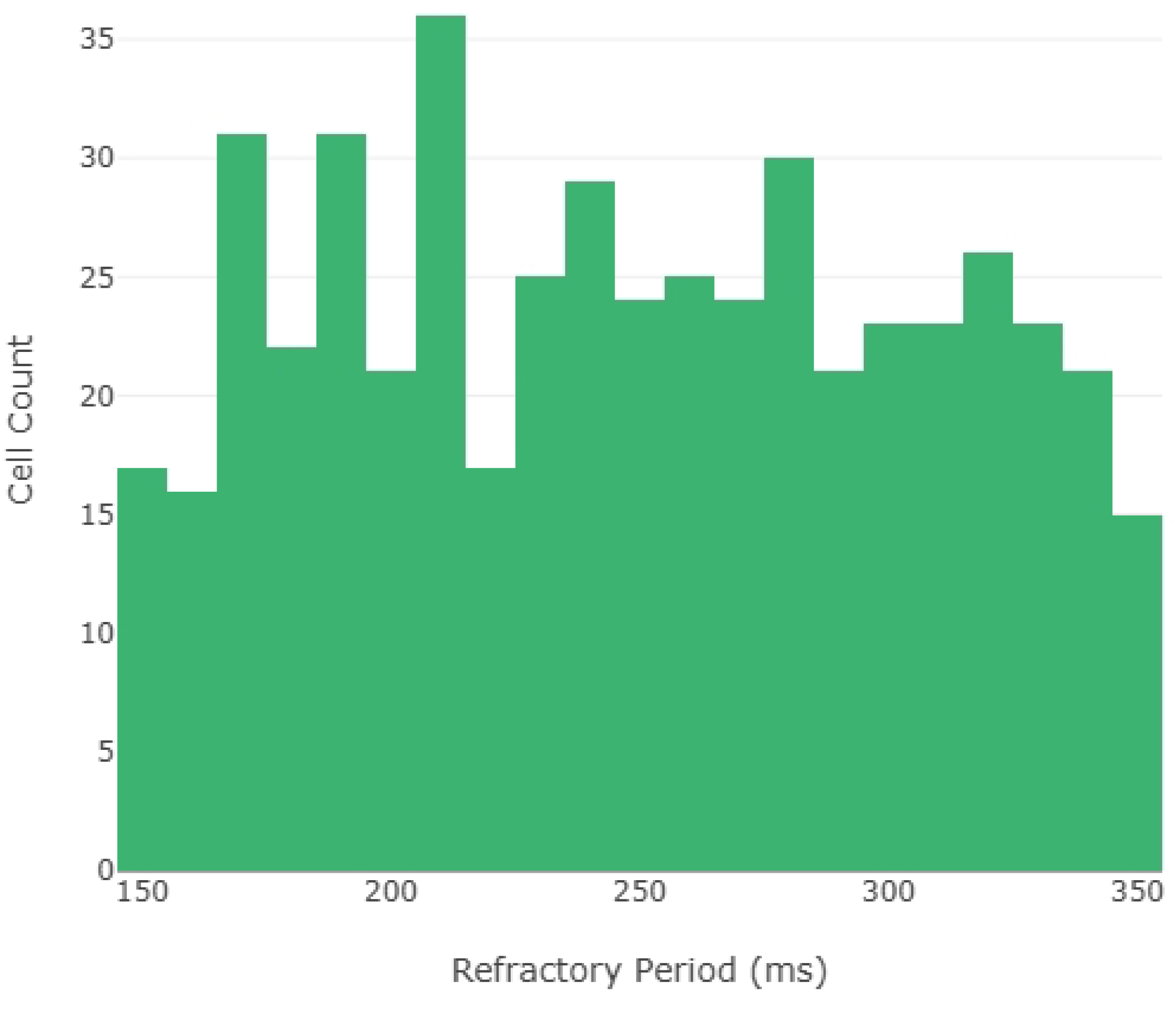
Histogram illustrating the distribution of conduction velocities assigned to the Purkinje cells. The variability in conduction speed reflects physiological heterogeneity, with values ranging from 0.3 to 2.0 m/s.

**Figure 1C:**
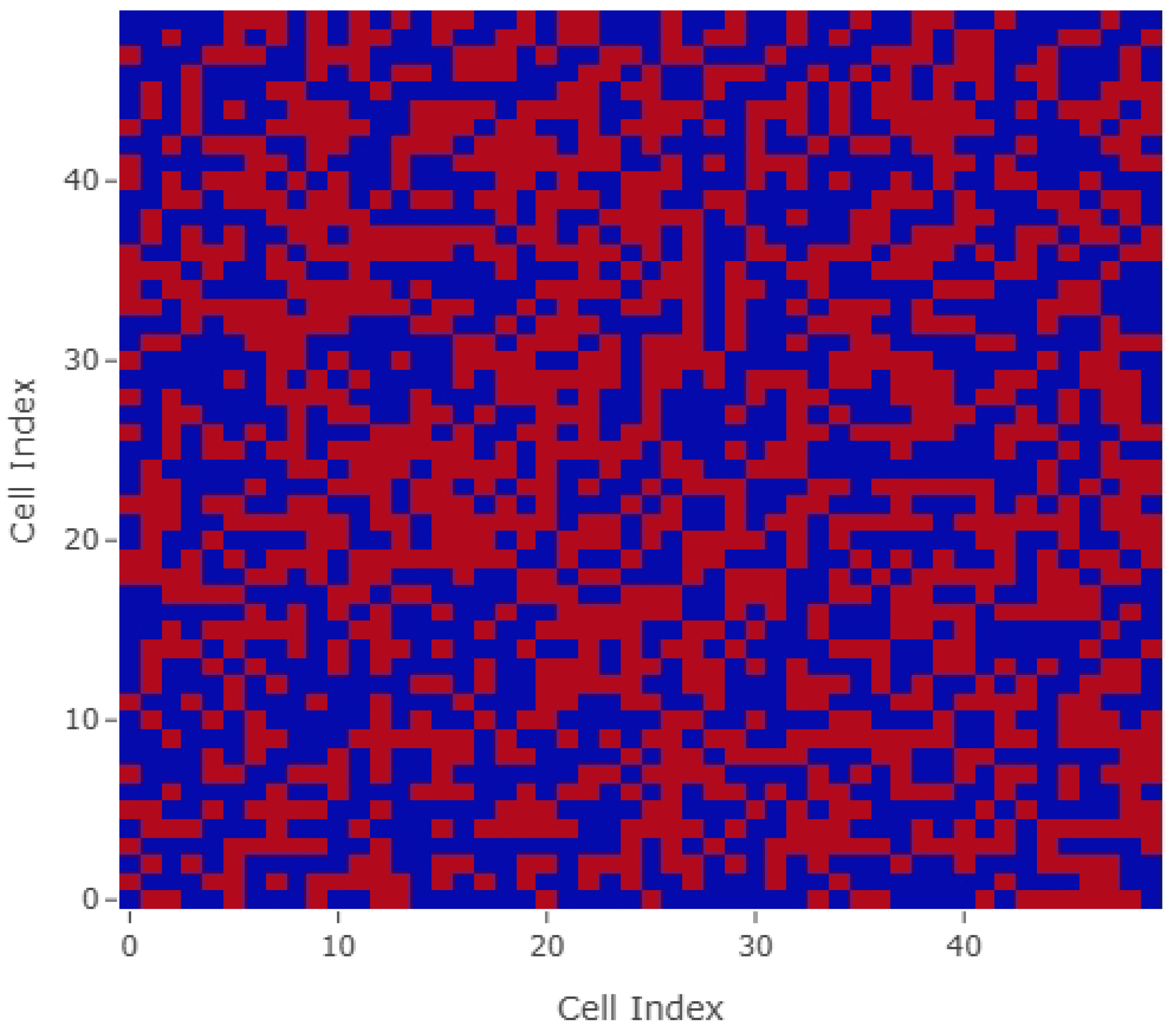
Visualization of ischemia severity gradients applied across two ischemic zones in the Purkinje network. One region demonstrates a linear severity decline, while the other applies an exponential decay gradient, simulating different ischemic injury patterns.

**Figure 1D:**
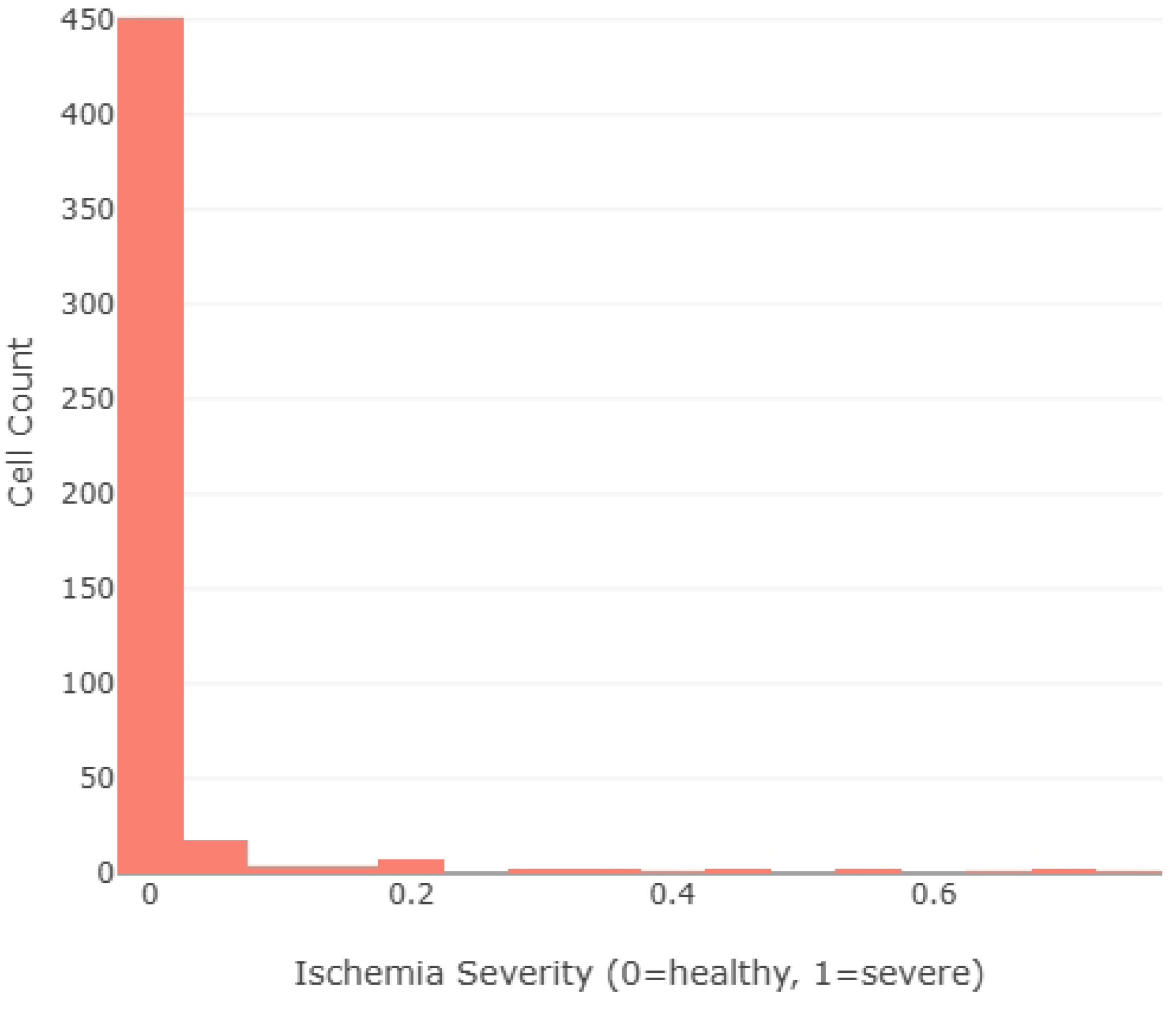
Heatmap representation of the network adjacency matrix showing binary connections between Purkinje cells. The randomized connectivity results in heterogeneous local coupling densities while maintaining overall network integrity.

**Figure 1E:**
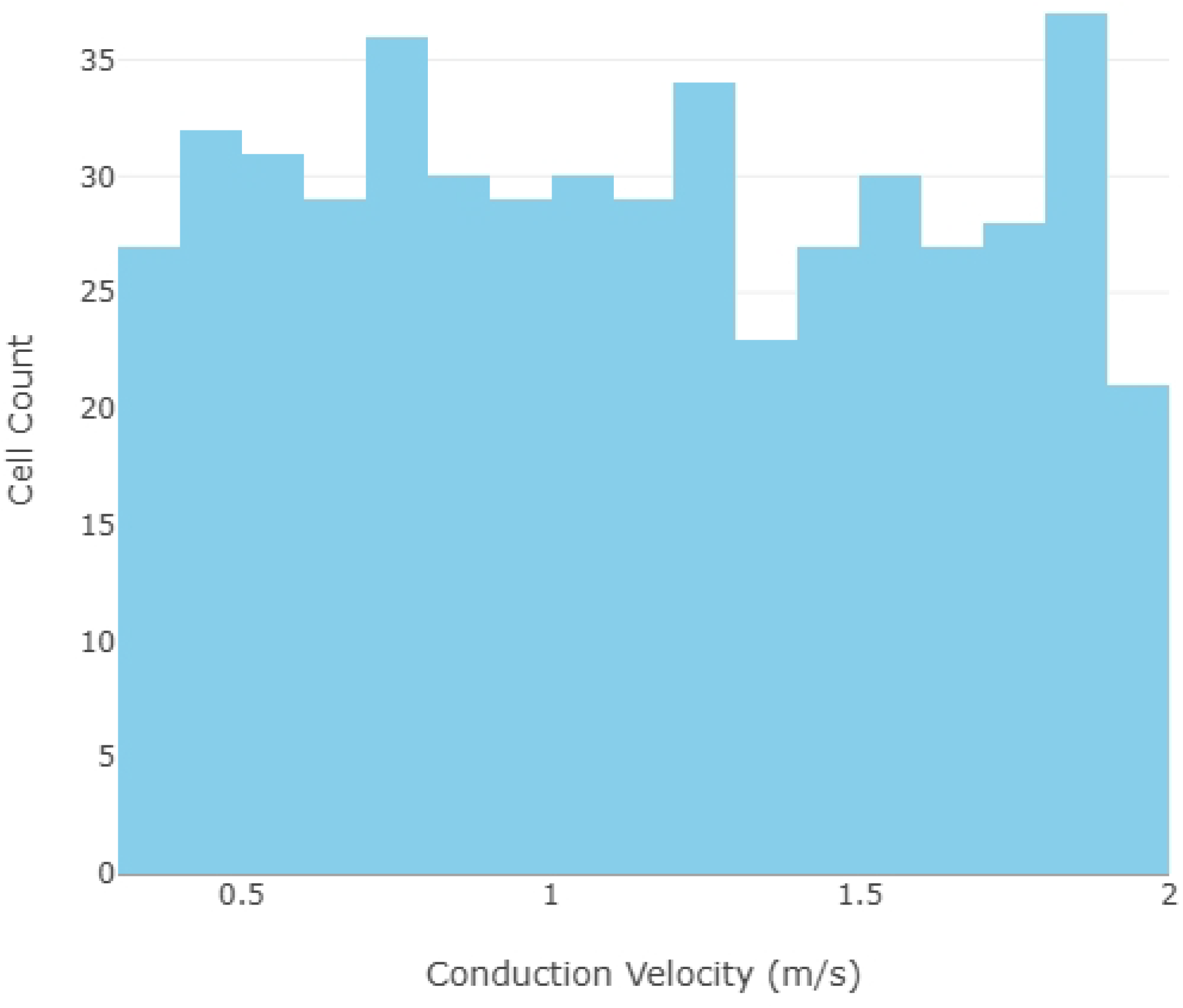
Histogram depicting the distribution of refractory periods assigned to individual Purkinje cells, ranging between 150 and 350 milliseconds, consistent with experimentally observed values.

**Figure 1F:**
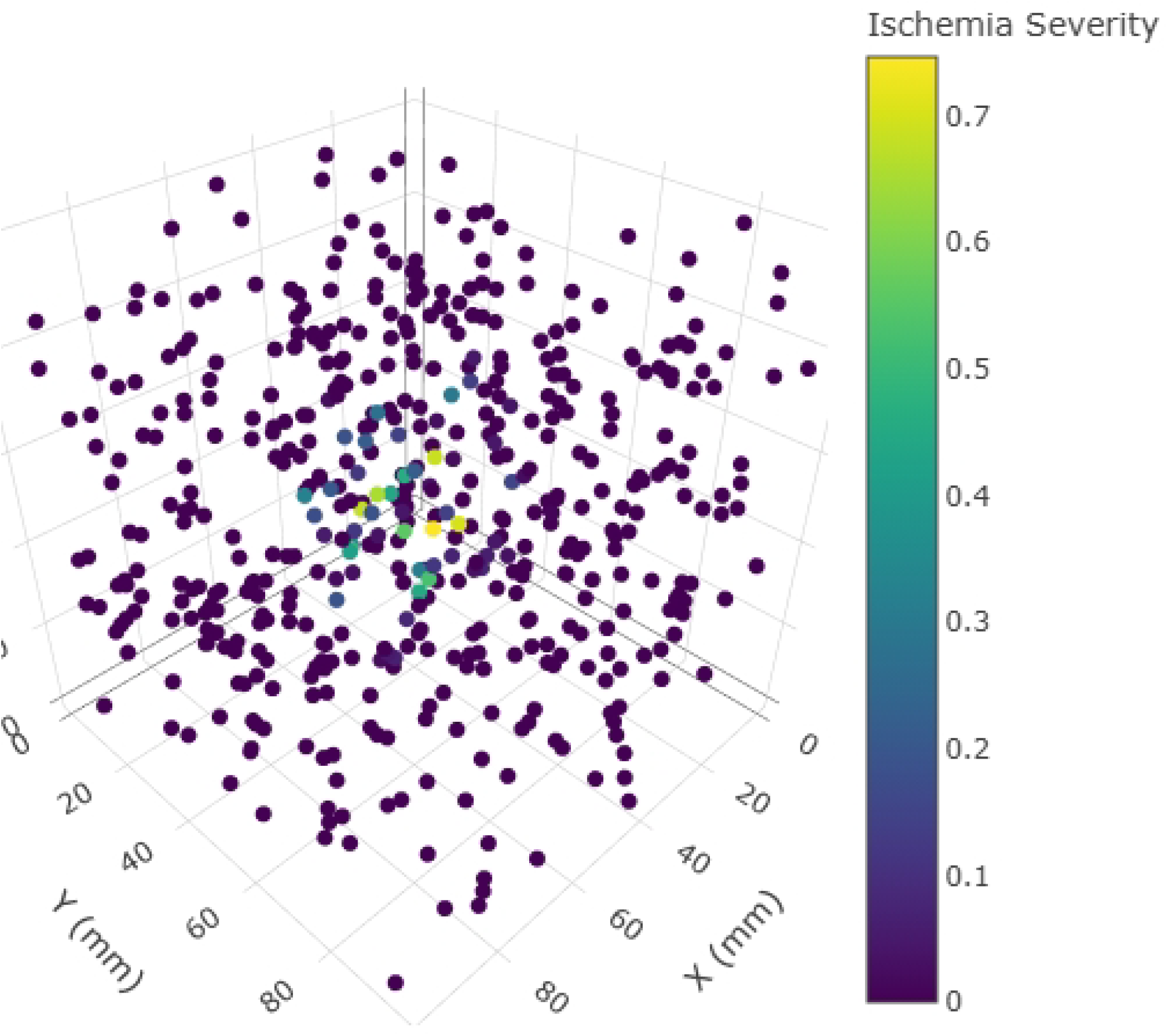
Two-dimensional XY projection of the 3D network, providing an alternative view of spatial clustering, network density, and ischemic region overlap for enhanced visual interpretation.

